# A global phylogenetic regionalisation of vascular plants reveals a deep split between Gondwanan and Laurasian biotas

**DOI:** 10.1101/2021.04.09.439199

**Authors:** Angelino Carta, Lorenzo Peruzzi, Santiago Ramírez-Barahona

## Abstract

Existing global regionalisation schemes for plants consider the compositional affinities among biotas, but these have not considered phylogenetic information explicitly. Incorporating phylogenetic information may substantially advance our understanding of the relationships among regions and the synopsis of biogeographical schemes, because phylogeny captures information on the evolutionary history of taxa. Here, we present the first phytogeographical delineation of the global vascular flora based on the evolutionary relationships of species.

We analysed 8,738,520 geographical occurrence records for vascular plant species together with a time-calibrated phylogenetic tree including 67,420 species. We estimated species composition within 200 × 200 km grid cells across the world, and used a metric of phylogenetic beta diversity to generate a phylogenetic delineation of floristic regions. We contrasted these results with a regionalisation generated using a taxonomic beta diversity metric.

We identified 16 phylogenetically distinct phytogeographical units, deeply split into two main clusters that broadly correspond to the Laurasia-Gondwana separation. Our regionalisation broadly matches currently recognized phytogeographical classifications, but also highlights that the Gondwanan area is split into a large Holotropical cluster and an Australian-NeoZelandic-Patagonian cluster. In turn, we found that the northernmost and southernmost units have the most evolutionarily distinct vascular plant assemblages. In contrast, taxonomic dissimilarity returned a regionalisation composed of 23 units with a high degree of shared taxa between Laurasian and Gondwanan areas, with no clear split among their biotas.

The integration of phylogenetic information provided new insights into the historical relationships among phytogeographical units, enabling the identification of three large, clearly differentiated biotas, here referred to as kingdoms: Holarctic, Holotropical, and Austral. Our regionalization scheme provides further evidence for delineating transition zones between the Holarctic and Holotropical kingdoms. The latitudinal patterns of evolutionary distinctiveness of vascular plant assemblages are consistent with recent evidence of higher and more recent diversification of flowering plants outside tropical regions.

## Introduction

Each species has a unique geographic distribution, but many species show similar geographic ranges. Shared ranges between two (or more) species may be due to a common evolutionary history, physical barriers to dispersal, or ecological requirements that limit survival (Lomolino *et al.*, 2004; Posadas *et al.*, 2006). In many cases, this translates into closely related species being distributed within the same regions more often than expected by chance, which in turn results in distinct lineages having non-random and spatially clustered distribution ranges (Lomolino *et al.*, 2004). As a result, different regions of the globe host different sets of living organisms (biotas).

These non-random geographic patterns are the prerequisite for dividing the earth into distinct biogeographical units (De Candolle, 1820), which should also take into account the phylogenetic relationships among taxa (Wallace, 1876). A biogeographical regionalisation is typically seen as a hierarchical system that classifies geographic areas according to their shared biotic composition. These hierarchical relations implicitly represent a shared evolutionary history among areas (Daru *et al.*, 2017). Thus, biogeographical regionalisation is the underlying framework for many basic and applied issues in ecology, evolution, and conservation (Kreft & Jetz, 2010; Morrone, 2018; Wen *et al.*, 2013). In principle, a biogeographical regionalisation should be constructed based not only on the distribution of species, but also on their phylogenetic relationships. Whilst biogeographers have generally used data available for the contemporary species distributions (Good 1947; Takhtajan, 1978; Cox, 2001), these analyses have been done without explicitly considering phylogenetic relationships among species, mostly due to the limited availability of large-scale phylogenies. In recent decades, species-level phylogenies based on molecular data have become increasingly available (*e.g.*, Jetz *et al.*, 2012; Jetz & Pyron, 2018; Smith & Brown, 2018; Upham *et al.*, 2019), allowing a more robust delimitation of biogeographic units based on historical relationships (Ronquist & Sanmartín, 2011; Holt *et al.*, 2013; Daru *et al.*, 2016; Ye *et al.*, 2019; Pataro *et al.*, 2020; Qian *et al.*, 2020).

Whilst global zoogeographical regions have been recently updated based on modern approaches (Holt *et al.*, 2013), the most recent global proposals for plants heavily rely on the scheme advanced by Cox (2001), which in turn is a qualitative refining of the floristic kingdoms proposed by Takhtajan (1978, 1986). Several proposals of a global phytogeographical regionalisation have been advanced to recognize between three and six floristic kingdoms (*sensu* Good 1947; Table 1). However, all these proposals have relied on qualitative criteria for the recognition of floristic kingdoms, and none has used explicit phylogenetic information in their biogeographical schemes. Recently, Procheş & Ramdhani (2020) attempted a global phytogeographical regionalisation based on the distribution and endemism of ancient plant lineages across the 35 phytogeographic units delimited by Good (1974). However, phylogenetic information (based on Harris & Davies, 2016) is only used for the identification of taxonomic units (lineages), but is not incorporated into the establishment of relationships among phytogeographic units.

**Table 1.** Phytogeographic kingdoms (in bold) and their historical circumscriptions. To describe major phytogeographic units, Engler (1879, 1882) coined the term “Realms”, which was later changed to “Kingdoms” by Good (1947).

|  | Engler (1879, 1882) | Diels (1908) | Good (1947) | Takhtajan (1978) | Cox (2001) | Morrone (2015) | Our proposal |
| --- | --- | --- | --- | --- | --- | --- | --- |
| [temperate and cold regions of the northern hemisphere] | **Arcto-Tertiary** | **Arcto-Tertiary** | **Holarktis** | **Holarctic** | **Holarctic** | **Holarctic** | **Holarctic** |
| [old world tropics] | **Palaeotropical** | **Palaeotropical** | **Palaeotropis** | **Palaeotropical** | **African**<br>[Cape region here] | **Holotropical** | **Holotropical**<br>Palaeotropical subkingdom<br>[Cape region here] |
|  |  | [New Zealand here] |  |  | **Indo-Pacific** |  |  |
| [new world tropics] | **Neotropical** | **Neotropical** | **Neotropis** | **Neotropical** | **South American**<br>[subantarctic islands here] |  | Neotropical subkingdom |
| [temperate and cold regions of the southern hemisphere] | **Ancient Ocean** | **Antarctic** | **Antarktis** | **Antarctic** |  | **Austral** | **Austral**<br>Antarctic subkingdom |
|  |  |  |  | [New Zealand here] |  |  | [Patagonia, New Zealand and SE Australia here] |
|  |  | **Australian** | **Australis** | **Australian** | **Australian** |  | Australian subkingdom |
|  |  |  | [New Zealand here] |  | [New Zealand here] |  |  |
| [Cape region in S Africa] |  | **Cape** | **Capensis** | **Cape** |  |  |  |

In this context, an integration of zoogeography and phytogeography into a single biogeographic scheme is needed (Morrone, 2015). However, the global biogeographic regionalisation based on plant distribution has lagged behind those constructed based on animals. To fill this gap, we carry out a reassessment of the phytogeographic regionalisation of the world that, for the firsts time, is based on a quantitative and phylogenetically informed big-data approach. We take advantage of publicly available global occurrence databases and available species-level molecular phylogenies for vascular plants, to integrate over eight million occurrence records with a dated phylogeny for more than sixty thousand vascular plants across the world. We applied pairwise phylogenetic (pβ) and taxonomic (β) beta diversity metrics to delimit geographical areas, discover their relationships, and identify the hierarchical organisation of the regionalisation. We then estimate the evolutionarily distinctiveness of the floras enclosed within phytogeographical units and discuss differences between our proposed framework with respect to previous regionalisations.

## Materials and Methods

### Geographic occurrence data

We retrieved geographic occurrence data for vascular plants across the globe from the Global Biodiversity Information Facility by querying all records under ‘Tracheophyta’ (we only considered ‘Preserved Specimens’ in our search); this database consisted of 68,570,538 occurrence records (GBIF.org, 2019). The occurrence records were taxonomically homogenized and cleaned following the procedures described in Edwards et al. (2015) and Ramírez-Barahona et al. (2020), but using Kew’s Plants of the World database as the source of taxonomic information (POWO, 2019; last accessed June 2019). Basically, we discarded records without species identification, flagged records with missing or badly formated GPS coordinates, and identified records potentially associated with biodiversity institutions. We updated family and species names through cross-validation with Kew’s taxonomic database, and used the CoordinateCleaner package (Zizka *et al.*, 2019) in R (R Core Team 2020) to flag suspect occurrence records meeting one of the following criteria: (1) equal latitude and longitude; (2) zero latitude and longitude; (3) coordinates falling within a five kilometre radius of a country’s political centroid; (4) coordinates falling within ten a kilometre radius of a country’s capital; (5) coordinates falling within a two kilometre radius of biodiversity institutions; and (6) coordinates falling in the open ocean, using a reference landmass buffered by one degree from the coastline to avoid eliminating species living in the coast.

By this approach we retained 27,537,044 records (40% of the original records) for 277,597 species. However, after inspection of the resulting database, we still detected several problems with the distribution of species, mostly associated with possible non-native records. Thus, we performed a complementary step to flag suspect records using Kew’s Plants of the World database, which includes information on the native status for many species in countries across the world. We flagged an additional 11,783,185 records that potentially represent non-native species distributions. The final geographic occurrence dataset used consisted of 15,753,859 records (23% of the original records) for 268,425 species of lycophytes, monilophytes, gymnosperms, and angiosperms across the world.

We incorporated point locality information into species distribution ranges (representing the maximum geographical extent of each species) by modelling point occurrences using hull geometries following the procedure described in Rabosky et al. (2016). Species with ≥ 6 unique occurrences (52,649 species) were modelled with alpha hulls (Roll *et al.*, 2017). Parameters for the delimitation of alpha hulls were adaptively selected using the function “getDynamicAlphaHull” from the package rangeBuilder (https://github.com/ptitle/rangeBuilder) in R (R Core Team 2020). We started with initial alpha values of two and then adjusted for the distribution of records so that at least 95% of the records were included within the estimated range. Species with 3–5 unique occurrences (6,767 species) were modelled using convex hulls. For species with 1–2 unique occurrences (7,853 species), we used a 7,584 km buffer based on the average spatial error reported in GBIF, to account for potential measurement and spatial errors. Species occurrences were extracted over an equal-area grid (Mollweide projection) based on 200 × 200 km grid cells. Finally, we also excluded grid cells containing fewer than two recorded species (Kreft & Jetz, 2010; Ramírez-Barahona *et al.*, 2020). Only few records occurred in Antarctica, so this region was not included in our analyses.

### Phylogenetic data

Initially, we used the species-level phylogenetic tree published by Jin & Qian (2019) (‘GBOTB.extended’), that includes 74,531 species of vascular plants, but with a poor representation of ferns (only 486 species out of ~ 9,000). To fill this gap, we combined the ‘GBOTB.extended’ tree with the fern phylogeny published by Testo & Sundue (2016) (‘TS.full’), which encompasses 4,007 fern species. Prior to combining the trees, we performed a taxonomic homogenization of the names in the two trees using Kew’s Plants of the World database, resulting in: 1) 1,720 synonyms flagged in the ‘GBOTB.extended’ tree; and 2) eight synonyms flagged in the ‘TS.full’ tree, with an additional 76 species that were not found in Kew’s database. After dropping these species from the trees, we merged the two trees by replacing the fern clade in the ‘GBOTB.extended’ tree with the ‘TS.full’ tree (without the outgroup); however, due to differing age estimates for the crown group of ferns between the two trees, prior to merging, we rescaled the ‘TS.full’ tree using the crown age for ferns provided in the ‘GBOTB.extended’. Thus, the final combined phylogenetic tree (‘GBOTB_TS’) keeps the original divergence age estimates of the ‘GBOTB.extended’ tree. The ‘GBOTB_TS’ tree was used in subsequent analyses and encompasses 76,226 species of lycophytes, monilophytes, gymnosperms, and angiosperms.

The taxonomic homogenization of the ‘GBOTB_TS’ tree allowed us to directly match species in the phylogeny to the geographic occurrence database. In sum, we obtained geographic data for 67,269 species of vascular plants included in the phylogeny, representing 8,717,249 occurrence records across the world.

### Computing phylogenetic and taxonomic beta diversity

Phylogenetic dissimilarity matrices among grid cells were calculated with Simpson’s phylogenetic pairwise beta diversity metric (pβ_sim_). Simpson index has the advantage of being independent of differences in species richness observed among sites (Kreft & Jetz, 2010). Phylogenetic beta diversity quantifies the turnover in phylogenetic composition between adjacent grid cells; this metric is an extension of the taxonomic beta diversity metric (β_sim_), where the proportion of shared species is instead substituted for the proportion of phylogenetic branch lengths represented by species shared between grid-cells. We constructed a matrix of pairwise pβsim among grid-cells and, for comparison, we also constructed a matrix for the taxonomic beta diversity metric (β_sim_) among grid-cells. All analyses were conducted using the package phyloregion (Daru *et al.*, 2020) in R (R Core Team, 2020).

### Cluster algorithm selection

The optimal number of units was defined using the ‘elbow’ method, setting the maximum number of clusters to k = 30 (Daru *et al.*, 2020). The ‘elbow’ method identifies the optimal number of units based on the range of explained variances (Daru *et al.*, 2020). We additionally evaluated the sensitivity of the resulting regionalisation scheme by setting the number of units to match the number of previosly recognized floristic kingdoms: k = 3, 4, 5, 6. The phytogeographical units identified were mapped and visualised using a multidimensional scaling colour space, indicating the degree of phylogenetic (or taxonomic) differentiation between units: phytogeographical units with similar colours have a greater proportion of shared clades (or taxa) that those with diverging colours. We then used the function ‘phyloregion’ to estimate the evolutionary distinctiveness (ED) of each phytogeographical unit by computing the mean value of phylogenetic (or taxonomic) beta diversity between a focal unit and all other units (Holt *et al.*, 2013; Daru *et al.*, 2020). All the analyses were conducted using the pβsim and βsim separately, in order to delineate a taxonomic and a phylogenetic regionalisation.

Nonmetric Multidimensional Scaling (NMDS) ordination plots and hierarchical clustering were used to evaluate the relationships among biogeographical units. Unit nomenclature mostly follows Takhtajan (1978). To identify spatial clusters of units across the world, we used the function “select_linkage” from the phyloregion package. This function assesses the degree of data distortion with the cophenetic correlation coefficient, which is a measure of how a dendrogram preserves the pairwise distances between the original distance matrix, and has a value between 0 (poor correlation) and 1 (strong correlation). We tested eight hierarchical clustering algorithms on the pβ_sim_ and β_sim_ matrices: single linkage, complete linkage, unweighted pair-group method using arithmetic averages (UPGMA), unweighted pair-group method using centroids (UPGMC), weighted pair-group method using arithmetic averages (WPGMA), weighted pair-group method using centroids (WPGMC), and Ward’s minimum variance. UPGMA was identified as the best clustering algorithm for both matrices (cophenetic correlation r = 0.66 for pβ_sim_ and r = 0.807 for βsim; Table S1).

## Results

We identified 16 phylogenetically distinct phytogeographical units across the world, which according to the hierarchical dendrogram of phylogenetic relationships are deeply split into two principal clusters that broadly match the separation between Gondwana and Laurasia (Fig. 1). The separation into these two clusters accounts for almost 50% of the total explained variance (0.31 out of 0.67; Fig. 1c). The hierarchical dendrogram also reveals that the Gondwanan cluster is divided into two subordinate clusters, albeit separated by lower pβsim values (Fig. 1d): one well defined Holotropical cluster (#2–9, 16), and a second Austral cluster composed of Australia (#7, 10) and a Neozelandic–Patagonian unit (#11). Indeed, the NMDS ordination (Fig. 1b) suggests that the Neozelandic-Patagonian and the Eremaean (#10) units are clearly isolated, but that Northern Australia (#7) has a closer affinity to some tropical units, namely the Malesian (#8), the Indian-Indochinese (#9) and, to a lesser extent, the Madagascan (#6) units.

**Figure 1.**
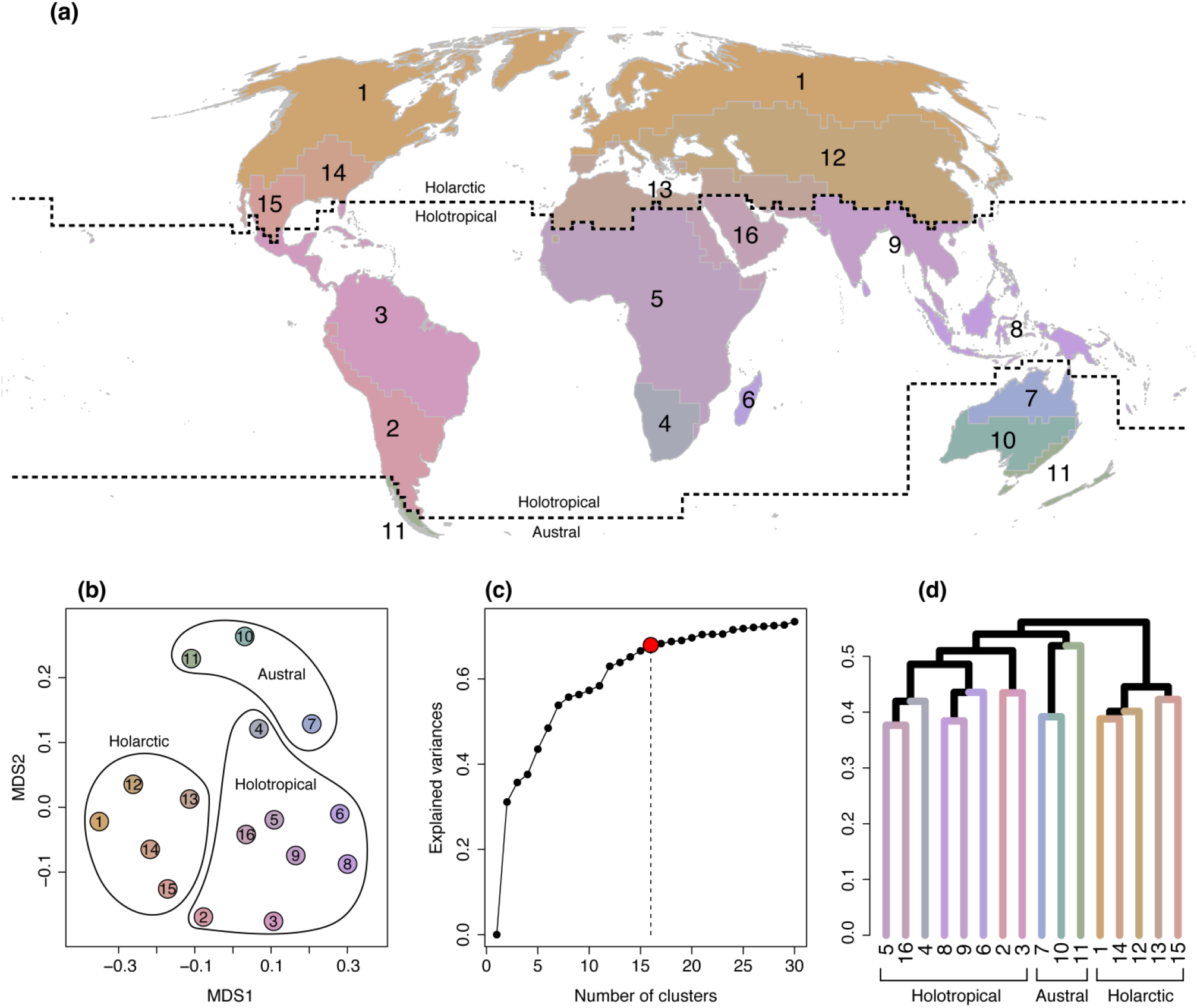
Map of the terrestrial phylogenetically distinct phytogeographic regions of the world (a), and their relationship presented as an NMDS ordination plot (b) and as a UPGMA dendrogram (d) of beta diversity (pβ_sim_) estimated across 200 × 200 km grid cells. Sixteen units were defined based on the ‘elbow’ method, considering the range of explained variance (c). Colours differentiating between units in the NMDS plot, dendrogram and map are identical. Dashed lines highlight the borders of the three kingdoms: Holarctic, Holotropical, and Austral, corresponding to the three major clusters in (d). NMDS stress = 0.156.

The Holotropical cluster is further split into two clusters separating the Neotropics and the Paleotropics: one includes most of Southern America (excluding Patagonia), Mesoamerica and the Antilles (#2, 3), and the second includes most of Africa (excluding North Africa), Southeast Asia, India, Malesia, and the Arabian pensinsula (#4, 5, 6, 8, 9, 16). The Madagascan (#6) unit, which is the smallest (Table 2), is phylogenetically more similar to the Malesian (#8) and the Indian-Indochinese (#9) units than to continental Africa (Fig. 1b-c). Lastly, the Laurasian cluster exhibits a higher internal homogeneity and harbours the largest phytogeographical unit, the Circumboreal unit (#1, Table 2). Here, the Madrean unit (#15) and the Mediterranean-Iranian unit (#13) cluster together.

**Table 2.** Summary of phytogeographic regions within each kingdom and subkingdom based on UPGMA clustering of phylogenetic beta diversity (pβ_sim_) for assemblages of vascular plant species within 200 × 200 km grid cells across the world. PD = total phylogenetic diversity; SR = species richness; ED = evolutionary distinctiveness.

| Kingdom | Subkingdom | Region | Unit# | Area in<br>km<sup>2</sup> | Total PD | SR | ED |
| --- | --- | --- | --- | --- | --- | --- | --- |
| **Holarctic** |  |  |  |  |  |  |  |
|  |  | Circumboreal | 1 | 50,920 | 113,364 | 11,058 | 0.561 |
|  |  | Eurasiatic | 12 | 22,520 | 146,542 | 14,870 | 0.534 |
|  |  | Mediterranean-Iranian | 13 | 10,760 | 72,027 | 6,867 | 0.498 |
|  |  | North-American-Atlantic | 14 | 3,520 | 56,317 | 3,292 | 0.518 |
|  |  | Madrean | 15 | 2,840 | 78,633 | 5,907 | 0.517 |
| **Holotropical** |  |  |  |  |  |  |  |
|  | Neotropical |  |  |  |  |  |  |
|  |  | Andean-Argentinian | 2 | 7,240 | 112,298 | 9,367 | 0.518 |
|  |  | Neotropical | 3 | 20,480 | 168,800 | 17,878 | 0.527 |
|  | Palaeotropical |  |  |  |  |  |  |
|  |  | Southern African | 4 | 3,480 | 70,906 | 6,424 | 0.534 |
|  |  | African | 5 | 22,920 | 89,369 | 5,849 | 0.489 |
|  |  | Madagascan | 6 | 1,640 | 53,441 | 2,727 | 0.538 |
|  |  | Malesian | 8 | 12,680 | 103,389 | 5,942 | 0.546 |
|  |  | Indian-Indochinese | 9 | 9,200 | 138,093 | 10,010 | 0.513 |
|  |  | Arabian | 16 | 6,200 | 35,171 | 1,544 | 0.469 |
| **Austral** |  |  |  |  |  |  |  |
|  | Australian |  |  |  |  |  |  |
|  |  | Northern Australian | 7 | 4,480 | 65,267 | 3,166 | 0.526 |
|  |  | Eremaean | 10 | 4,640 | 61,400 | 3,785 | 0.551 |
|  | Antarctic |  |  |  |  |  |  |
|  |  | Neozelandic-Patagonian | 11 | 3,680 | 65,911 | 3,774 | 0.549 |

Even when reducing the number of units to match the number of currently recognized floristic kingdoms (k = 3, 4, 5, 6), the separation between the Gondwanan and Laurasian clusters is always maintained, as well as the independency of the Austral cluster, which includes Australia and the Neozelandic-Patagonian unit (Fig. S1-S4). For instance, for k = 6, the three main clusters (each with two units) are identifiable: Holotropical, Austral, and Holarctic (Fig. S4). In this case, the Holarctic cluster is composed of a large Circumboreal unit and of a Trans-Atlantic unit roughly corresponding to the transition zones between the northern and southern hemispheres. In turn, with k = 5 (Fig. S3), the Holotropical cluster is composed of a Indo-Malesian unit and a large amphi-Atlantic unit composed by Africa and the Neotropics, whereas the Austral cluster is divided into the Autralian and NeoZelandic-Patagonian units.

As expected, the pβ_sim_ and the β_sim_ values are strongly correlated (Mantel test: *r* = 0.68, *P* = 0.001, 999 permutations). However, the best taxonomic regionalisation is composed of 23 units (Fig. S5) and, despite the higher number of units, the total explained variance (0.5) is lower than that explained by the phylogenetic regionalisation presented above (0.67). In addition, when omitting phylogenetic information, we found a less clear phytogeographic distinction between the Gondwanan and Laurasian clusters. More specifically, in the taxonomic regionalization, the Mediterranean-Iranian unit (which is here split into two separate units) groups with the Paleotropical units, whereas the Madrean unit, together with an Appalachian and a Californian unit, is recovered within a mostly Neotropical cluster (Fig. S5). In addition, the Patagonian and Neozelandic units are not grouped together, with the former being placed within a Neotropical cluster and the latter placed within a Paleotropical cluster.

Overall, based on our phylogenetic regionalization, we show that the Northernmost and Southernmost units, together with the Malesian and Neotropical units, exhibit the highest evolutionary distinctiveness (Fig. 2a). Also, regions (#2, 7, 13, 15, 16) at the contact between main clusters generally show lower ED values.Since lycophytes, monilophytes, and gymnosperms could potentially have a strong impact on ED estimates because they represent less diverse, evolutionarily distinct clades, we re-ran our analysis considering angiosperms only. This resulted in similar ED patterns (Fig. 2b), but with overall higher ED estimates for angiosperms (Fig. 2b).

**Figure 2.**
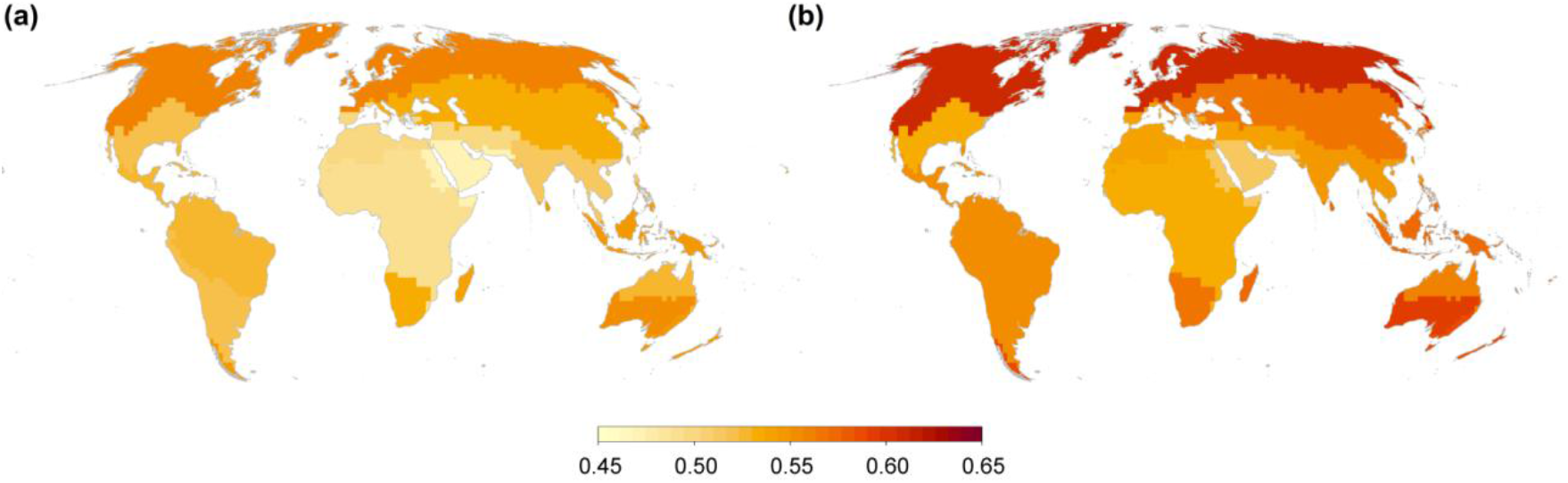
Evolutionary distinctiveness within the 16 phytogeographical units, considering all vascular plant species (a) and angiosperms only (b), quantified as the mean of pairwise pβ_sim_ values between each unit, contrasted with all other units. Darker regions indicate regions of higher evolutionary distinctiveness.

## Discussion

Here we built a comprehensive biogeographic regionalisation scheme of the world based on the integration of large-scale geographic and phylogenetic data for vascular plants. This allowed us to quantitively test, for the first time, century-old regionalisation of the world based on plant distributions (Engler, 1882; Diels, 1908; Good, 1947; Takhtajan, 1978). This is an important step forward in building a comprehensive, phylogenetically informed regionalisation of the world (Morrone, 2015). Integrating phylogenetic and occurrence records on a global scale, we were able to retrieve three major clusters, which we identify here as kingdoms, and their cladistic relationships following the sequence: (Holarctic, (Holotropical, Austral)). Both Holotropical and Austral kingdoms are split into two main clusters, which we identify here as subkingdoms, while the 16 main phytogeographical units which best explain the variance observed in our dataset are here referred as regions (Table 2). Because our goal is to identify major clusters of phytogeographical units and their relationships at a global scale, the hierachical organisation of our regionalisation is described using only higher rank categories (kingdom, subkingdom, region),whereas other lower-rank categories (*e.g.*, dominion, province, district) (Morrone, 2018) were not considered.

Our phylogenetic regionalisation of the world’s vascular flora into distinct phytogeographical kingdoms, subkingdoms, and regions show substantial congruence to long-recognised regionalisation systems (Engler, 1882; Diels, 1908; Good, 1947; Takhtajan, 1978; Cox, 2001; Morrone, 2015), including a deep separation of biotas broadly corresponding to the Laurasian-Gondwanan divergence (Raven & Axelrod, 1974; Morrone 2015). More specifically, our results are in agreement with the recent “integrative” biogeographical proposal advanced by Morrone (2015), in recognising just three major biogeographically unique areas of the world: Holarctic, Holotropical, and Austral. Our results recovered the general features of the regionalisation of Morrone (2015), but with some exceptions. The Holarctic Kingdom herein recognized is not clearly segregated into Nearctic and Palearctic components. Instead, this kingdom is split into two clusters: a) Circumboreal; and b) temperate Asia and the Mediterranean. Repeated migrations between Eurasia and North America (Donoghue & Smith, 2004; Graham, 2018) likely explain the homogeneity in phylogenetic composition of the Holarctic cluster, while keeping a high evolutionary distinctiveness. The Austral kingdom coincides with that recognized by Morrone (2015) and by Procheş & Ramdhani (2020), except for the Southern African region (linked to the Holotropical kingdom here) and the Antarctic territories (not assessed here). In turn, the Holotropical kingdom mostly corresponds to that recognized by Morrone (2015), but with a main split between the Neotropical and Palaeotropical subkingdoms, in agreement with most of classical phytogeographical proposals (Table 1). The split between Neotropics and Paleotropics has also been supported in a recent assessment of area relationships across the tropics (Slik *et al.*, 2018).

Although our results broadly match traditional regionalisation schemes, we also identified some discrepancies. For example, the phytogeographical relationships of New Zealand have long been debated (Moreira-Muñoz, 2007), being alternatively placed within either Palaeotropical, Australian, or Antarctic kingdoms (Table 1). Here, we clearly found a phylogenetically homogeneous unit, composed of New Zealand and Patagonia clustering with Australia, highlighting a common evolutionary history and isolation of these territories (Procheş & Ramdhani 2020). As proposed by Gonzalez-Orozco et al. (2014), Australia is separated into two regions that broadly correspond with tropical (north) and extratropical (south) Australia, the former showing a closer affinity with the Malesian region (Fig. 1). Although the South African region is clearly distinct from the rest of Africa (Fig. 1), this region is not recognised as a major unit (kingdom) as advocated by Diels (1908), Good (1947), and Takhtajan (1978), but instead it is nested within a broader African group; this is in agreement with Cox (2001), who did not recognise a Cape kingdom. Interestingly, the Southern African region is more clearly differentiated from the rest of Africa when only taxonomic turnover is considered (Fig. S5). In line with Takhtajan (1978), we also recognize a well distinct Arabian region (Brenan, 1978). Despite the close geographical proximity of Madagascar to Africa, and their taxonomic affinity (Fig. S5), the former exhibits a remarkably higher phylogenetic affinity with the Indo-Chinese, Malesian, and Australian regions, as already highlighted by Schatz (1996). The discrepancies between our results and traditional regionalisation schemes highlight the importance of explicitly incorporating phylogenetic information to establish phytogeographical schemes. This is further supported by the observed differences between the phylogenetically informed regionalisation and that based only on taxonomic turnover.

Our results also support an intermadiate position of some regions between kingdoms (Fig. 1b). More specifically, we found an intermediate position between Holarctic and Holotropical kingdoms of the Mediterranean and Madrean regions, which also exhibit an overall lower evolutionary distinctiveness (Fig. 2; Table 2). Based on our results, we support a view of these regions as transition zones, which result from the intergrading of independent evolutionary histories (Morrone, 2015). Usually, clustering algorithms are unable to differentiate between core and transition zones, because the former are typically merged with core areas with which they show the greatest affinity (Kreft & Jetz, 2010). Despite this, however, we found additional evidence for these transition zones from the hierarchical relationships: the Madrean and the Mediterraneo-Iranian regions (#15, 13) were retrieved as clustering separately from the rest of the Holarctic kingdom. These results support the hypotheses that taxa from these transitions zones are, in general, more closely related to Holarctic taxa than to taxa from tropical regions (Morrone 2015).

The contrast between the phylogenetically informed regionalisation and that based only on taxonomic affinities, demonstrates the advantages of incorporating phylogenetic information for the delineation of phytogeographical regions on a global scale. The phylogenetically informed regionalisation mostly support the proposed schematic diagram of historical area relationships proposed by Morrone (2015), which is mostly based on Engler (1982). Furthermore, the taxonomic-only approach is less parsimonious, and explains a lower amount of variance in phytogeographic affinities among units, resulting in more shallow differences among major clusters (Fig S5). In terms of the inferred relationships among units, one of the main differences between the phylogenetically informed and the taxonomic-only approaches concerns the northern limit of the Holotropical kingdom. Using taxonomic data only, the Holotropical kingdom also includes the Mediterranean–Iranian and the Madrean regions, further confirming the view of these units as transition zones. Another important difference between the two approaches is that the taxonomic-only regionalisation recognizes three alternative main clusters: Neotropical, Paleotropical, and Laurasian. Here the Laurasian cluster mostly corresponds to the Holarctic kingdom. However, in this case, the units corresponding to the Austral kingdom (otherwise recognized as a distinct cluster, Fig. 1) are split up into two separate areas: the first (Patagonian unit) is identified within the Neotropical cluster, and the second (Neozelandic unit) within the Paleotropical cluster. This regionalisation is less parsimonious than recognizing a single Austral kingdom, partly due to the high phylogenetic distinctiveness of its flora (Fig. 2). More specifically, the higher phylogenetic uniqueness of the Neozelandic-Patagonian region is consistent with the isolation of these areas and their connection through Antarctica, resulting in the presence of plant lineages (*e.g.*, Berberidopsidales, Araucariaceae) with disjunct distributions (Winkworth *et al.*, 2015; Procheş & Ramdhani 2020).

The observed latitudinal patterns of evolutionary distinctiveness are suggestive of phylogenetic niche conservatism (Wiens, 2004) likely leading to the accumulation of phylogenetic diversity within tropical regions, particularly for angiosperms and ferns (Schuettpelz & Pryer 2009; Ramírez-Barahona *et al.*, 2020). However, the latitudinal patterns of evolutionary distinctivenes are higher for angiosperms, and this can be interpreted in light of the presumed tropical origin of this group (Feild *et al.*, 2009; Coiro *et al.*, 2019) and of the delayed, but accelerated, rise of flowering plant lineages in extratropical regions (Igea & Tanentzap, 2020; Ramírez-Barahona *et al.*, 2020). Nonetheless, phylogenetic dissimilarity between the three kingdoms is not high, and instead is higher among phytogeographical regions. One possible biological interpretation is that the world’s flora shares a recent common evolutionary origin, especially for the overwhelmingly larger angiosperm component, and that dispersal across the world has played a key role in determining the distribution of plant lineages. This does not allow for stronger divergences among the three kingdoms, albeit these differences are sufficiently clear to recognise them as distinct biogeographical units.

The inclusion of phylogenetic information allowed a robust regionalisation of vascular plant distributions at a global level and provided new insights into the historical relationships among phytogeographical regions. The match between our regionalisation and earlier schemes is remarkable. Thus, we propose only some minor changes to the existing classification schemes, mainly at higher ranks. Our results could be a launch pad for further detailed studies, specifically devoted identifying and circumscribing lower rank units (i.e. below region). The phytogeographic regionalization herein presented provides a baseline for future ecological, evolutionary, and conservation studies of vascular plants at global scale and represents, together with the zoogeographical scheme advanced by Holt *et al.* (2013),, a major step forward towards the integration of zoogeography and phytogeography into a single biogeographical scheme.

### Data limitations and caveats

In any biogeographical analysis, acknowledging the limitations of the geographic and phylogenetic data is fundamental to properly interpret the resulting geographical patterns. Here, some limitations of the occurrence data include the long-recognized problems associated with publicly available data (*e.g.*, misidentification of species, erroneous geographic records, non-native records). The series of taxonomical and geographical filters we applied to the data were aimed at minimizing these problems. However, a main limitation in the data still remains: the geographical bias in occurrence records, which is evident at first sight in regions such as the Indian subcontinent, and Siberia, which are well-known to be considerably underrepresented in the GBIF database, and might lead to biased results. However, we believe that the recognition of the three main biogeographic kingdoms is robust enough to these sources of bias, but these could potentially have an impact on the delimitation (and especially relationships) among individual phytogeographical regions.

Phylogenetic data also comes with some limitations due to 1) incompletely sampled trees, 2) sampling bias across lineages, 3) topological resolution, and 4) uncertainty in divergence time estimates. The first two limitations probably have a minor impact on the results, given that the sampling is skewed towards the larger and more widespread lineages, yet some major lineages are poorly sampled in the phylogentic tree, such as the lycophytes and some monilophyte lineages (*e.g.*, tree ferns). The third limitation represents a more serious challenge to any proposed regionalisation, but it would only seriously affect the shallower relationships between species and genera, therefore being of relevance only for the delimitation of more nested areas within the kingdoms. The fourth is a major, but often neglected limitation in biogeographic analyses, where divergence time estimates are accepted at face value without consideration of uncertainties stemming from model assumptions and fossil calibration schemes (Pranham *et al.*, 2012; Sauquet *et al.*, 2012; Magallón 2021). In this context, the phylogenies used here (after rescaling of the fern phylogeny) proved to be useful to address the phytoregionalisation of the world, but differences in divergence times will most likely have a strong impact in the estimate of phylogenetic dissimilarity. These sources of uncertainty would play a major role when attempting to interpret any proposed regionalisation in terms of the evolutionary history of species.

## Data availability statement

The geographic occurrence data for vascular plants is available from the Global Biodiversity Information Facility with the identifier https://doi.org/10.15468/dl.bdxzkw.

## Author contributions

A.C. and L.P. conceived the idea; A.C. designed the research; S.R.B. compiled and curated the occurrence and phylogenetic data; A.C. performed the analysis; A.C. wrote the manuscript with contributions from L.P. and S.R.B.

